## Supplementary Figures for "Melanization of *Candida auris* is Associated with Alteration of Extracellular pH"

### $^{15}$N CPMAS ssNMR of *C. auris* cells non-melanized, CDC 388

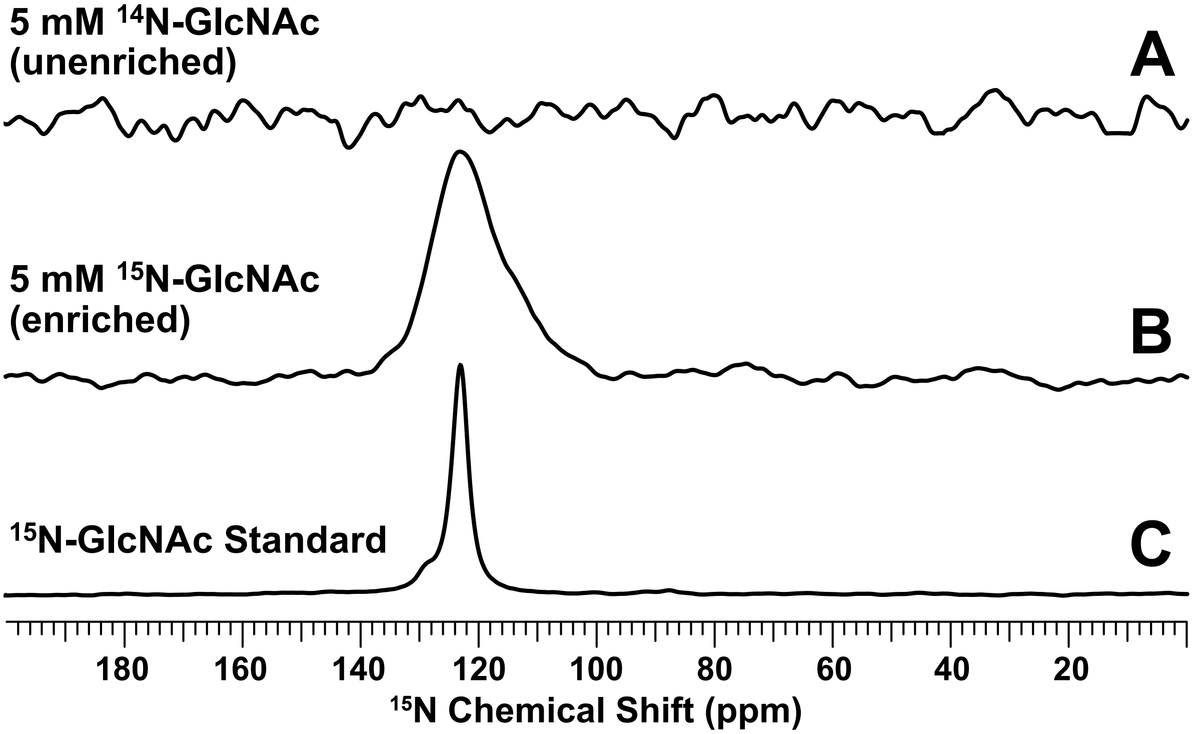


***Supplementary Figure 1. C. auris takes up exogenous GlcNAc and uses it for chitin synthesis.*** $^{15}$N cross-polarization solid-state NMR spectra of intact heat-killed *C. auris* cells from the melanizing strain CDC 388 grown with (A) 5 mM $^{14}$N-GlcNAc or (B) 5 mM $^{15}$N-GlcNAc; (C) spectrum of a $^{15}$N-GlcNAc standard compound. All data were acquired with 64 scans (~3 minute experimental time). The absence of signal in the spectrum of $^{14}$N-GlcNAc supplemented cells (A) after the acquisition of only 64 scans is consistent with the low natural abundance (0.4%) of the NMR-active $^{15}$N isotope. The presence of signal in the spectrum of $^{15}$N-GlcNAc supplemented cells (B) after the acquisition of only 64 scans is consistent with the uptake of substrate enriched in the $^{15}$N-isotope. The $^{15}$N chemical shift (~123 ppm) of the observed signal is consistent with the amide nitrogen within an acetyl group; its relatively broad linewidth in comparison to that of the peak in spectrum (C) is typical of large macromolecules, indicating that the taken-up exogenous GlcNAc was subsequently used for chitin synthesis.

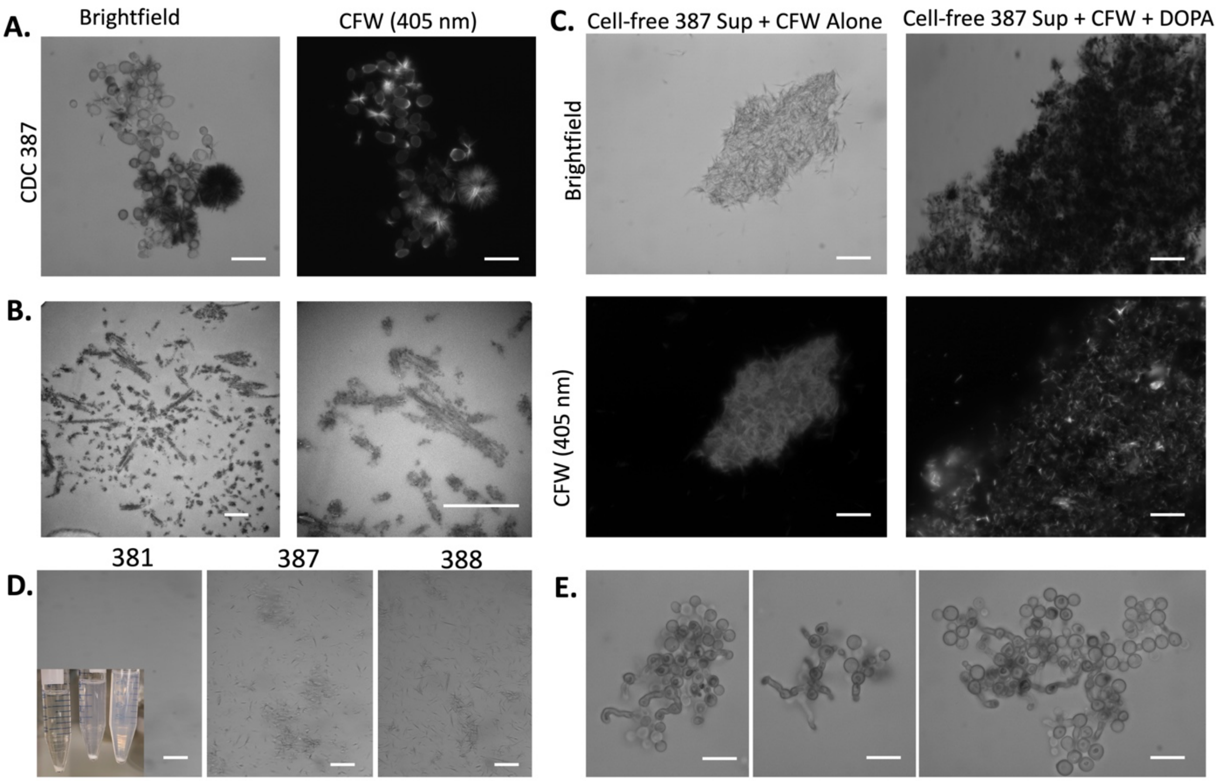


***Supplementary Figure 2. Interactions of Calcofluor White and melanin.*** (A) Melanizing *C. auris* strains (CDC 387) grown with L-DOPA and Calcofluor White (CFW) have visible crystal formation under light microscopy. These crystals are dark in color, suggesting the presence of melanin or polymerized L-DOPA, and they are fluorescent under a 405 nm wavelength laser, indicating they also contain CFW. (B) These crystals can also be seen with transmission electron microscopy. (C) Adding L-DOPA and CFW to non-melanized cell-free supernatant resulted in the formation of black crystals, which under the fluorescent microscope have signal, indicating that the crystals are a combination of both CFW and melanin/autopolymerized L-DOPA. (D) Additionally, when CFW is added to supernatant, the crystals that form in CDC 381 strain are large and settle quickly, leaving few crystals in suspension, whereas CDC 387 and 388 form smaller crystals that do not settle rapidly and remain in the suspension. Inset is an image of the falcon tube containing the supernatant with CFW, and micrographs show a sample taken from the non-settled suspension. (E) CDC 388 cells grown in 50% MIC Caspofungin (0.5 $\mu$g/ml) demonstrating pseudohyphal growth. Scale bars in A, C, D, and E represent 10 $\mu$m, and scale bars in B represent 500 nm.

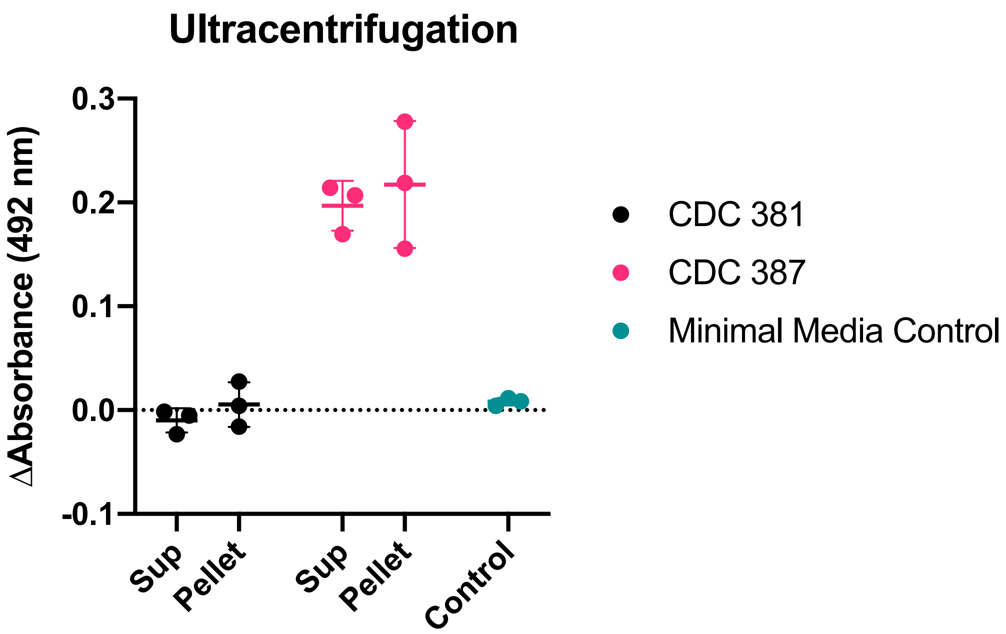


***Supplementary Figure 3. Melanization activity is not associated with extracellular vesicles.*** Following ultracentrifugation of the *C. auris*-conditioned media, there is comparable melanization activity in both the supernatant and the pellet (containing extracellular vesicles) resuspended in 200 $\mu$l of supernatant, which represents a ~200x concentration.
